# The rabies virus interferon antagonist P protein selectively modulates interferon signalling to inhibit antiviral gene expression while supporting proviral gene expression

**DOI:** 10.64898/2026.07.30.737726

**Authors:** Alebachew Messele Kebede, Cassandra T. David, Stephen M. Rawlinson, Celine Deffrasnes, Paul R. Gooley, Samuel C Forster, Gregory W. Moseley

## Abstract

Type-I IFNs mediate the principle antiviral response of cells by controlling the expression of hundreds of IFN-regulated genes (IRGs), many of which have antiviral functions. The best understood mediators of IFN signalling are STAT1 and STAT2, and STAT1/2-dependent gene induction is conventionally viewed as the primary outcome of type-I IFN signalling. To overcome the IFN response, viruses express proteins called IFN-antagonists, which target IFN signalling pathways (e.g. rabies virus P-protein (RABV-P) binds and inhibits IFN-activated STAT1/2) and so are typically considered to mediate shutdown of the IFN response. However, IFN signalling is not exclusively antiviral, with many IRGs reported to be required for or to facilitate infection by certain viruses. How viruses coordinate the apparent need to suppress certain antiviral IRGs, while presumably permitting the expression of others, including ‘proviral’ IRGs is poorly defined. However, it has been shown that type-I IFN can activate multiple pathways other than classical STAT1/2, so discriminatory targeting of specific pathways by IFN-antagonists may enable highly selective regulation of distinct IRGs, dependent on the requirements of the specific virus.

Here, we analyse the global effects of RABV-P protein on the IFN-regulated transcriptome. We confirm that IFN not only stimulates (IFN-stimulated genes, ISGs) but also represses (IFN-repressed genes, IRepGs) a large number of IRGs. Notably, our data indicate that RABV-P protein can antagonize both the IFN-dependent stimulation and repression of certain ISGs and IRepGs, without significantly impacting the expression of large proportion of IRGs. Antagonized ISGs included classical antiviral genes, while non-antagonized ISGs include genes with pro-viral effects on RABV. Transcription factor analysis indicated that RABV-P antagonizes STAT1/2-regulated IRGs, but not IRGs regulated by other pathways including MAP-kinase pathways, which are important to process such as cell survival and inflammatory response. These data indicate that selective modulation rather than global inhibition of IFN-signalling has beneficial outcomes for replication. The data also support the significance of IRepGs, and their modulation by IFN antagonists in viral infection.

**Significance Statement:** The ability of viruses to evade immunity is critical to disease and so presents targets for the development of interventions. The principle antiviral response of cells is mediated by interferons (IFNs), which activate STAT proteins to induce hundreds of genes including antiviral genes. Viruses counter this by expressing ‘IFN-antagonist’ proteins, many of which directly inhibit STATs. IFN-antagonists are typically considered to shut down IFN responses, but many IRGs have ‘pro-viral’ functions. How viruses coordinate the apparent need to antagonise some IRGs but not others are poorly understood. Using a well-characterised viral IFN-antagonist, we find that by selectively targeting certain IFN-activated pathways, IFN-antagonists can inhibit effects of IFN on specific subsets of IRGs (including antiviral genes) without affecting others (including proviral genes); thus, IFN-antagonists may be redefined as selective ‘IFN-modulators’.

## Introduction

In humans interferons (IFNs) comprise type-I, II and III, which differ in properties including structure and receptor distribution (1). The type-I IFN family comprises multiple subtypes including IFN-α (which has 13 subtypes) and IFN-β, which are the major mediators of the innate antiviral response (2). The first stage of the IFN response is the induction of IFN where, following the infection of cells and detection of pathogen associated molecular patterns (PAMPS, such as viral nucleic acids) by pathogen recognition receptors (PRRs), a signalling cascade is activated. This results in the activation by phosphorylation of transcription factors (TFs) including IFN-regulator factor 3 (IRF3) and, in certain cell types or conditions, IRF7 (3). IRF3 and IRF7 translocate to the nucleus and bind to promoter sites of IFN-α/β genes to activate IFN expression. IFN-α/β can be produced by most nucleated cell types in response to infection and forms the principal innate immune response of cells to viral infections (4). Following release from cells, IFN-α/β signal in autocrine and paracrine fashion to induce an antiviral state by binding to IFN-α/β cell surface receptors (IFNAR) and activating IFN-signalling (3).

Classical signalling downstream of the IFNAR complex is primarily mediated by the TFs signal transducers and activators of transcription 1 and 2 (STAT1/2). Specifically, ligation of IFNAR by IFN activates receptor associated Janus kinases (JAK1 and TYK2) which phosphorylate conserved tyrosines in STAT1 and STAT2 (5). The tyrosine-phosphorylated (pY) STATs form pY-STAT1/2 heterodimers via reciprocal interactions of the phosphor-tyrosine and Src homology 2 (SH2) domains of each STAT protein (6). The heterodimers translocate to the nucleus where, together with IRF9, they form the IFN-stimulated gene factor 3 (ISGF3) complex that binds to IFN-stimulated response element (ISRE)-containing promoters of IRGs, stimulating the expression of several hundred ISGs (2). ISGs mediate diverse effector functions including antiviral, antiproliferative, proapoptotic and immunomodulatory functions, which can result in the suppression of viral replication, and foster the development of an adaptive immune response (7, 8). IFN also suppresses the expression of a large number of genes (IFN-repressed genes, IRepGs)) (7). Type-I IFN regulated IRepGs are associated with functions including the regulation of cell growth, apoptosis and immune response (9, 10); however their specific role in antiviral response is less well defined. Emerging evidence suggests that type-I IFN IRepGs include host-dependency factors, which are cellular components that viruses hijack for infection/replication processes (11), such that repression of their expression may contribute to the IFN-induced antiviral state by ‘starving’ viruses of these required components. Thus, the type-I IFN response forms an essential component of host defence against viral infections, consistent with observations that the absence of an IFN response leads to increased susceptibility to various viral infections (3, 12, 13). Together, these data have established the view that signalling by type-I IFNs forms an antiviral system, primarily dependent on pY-STAT1/2 (3).

To counteract the type-I IFN response, viruses express proteins collectively referred to as IFN-antagonists (14). These proteins can inhibit the IFN system at multiple stages and by diverse mechanisms that can differ significantly between viruses (15, 16). The IFN-activated signalling pathway is a common target of IFN antagonists, through mechanisms including the expression of decoy IFN receptor proteins, or targeting of the IFNAR or downstream components (e.g. JAKS, STATs) through mechanisms including degradation, inhibition of phosphorylation, or mislocalization (14, 17, 18). The rabies virus (RABV) P-protein (RABV-P) is one of the most extensively characterised IFN-antagonists, and can target the induction, signalling and effector stages of the IFN response. In IFN induction, RABV suppresses IRF3 phosphorylation and subsequent expression of type-I IFNs (19, 20). In the IFN signalling stage, RABV-P protein interacts directly with pY-STAT1-containing complexes (including homodimers with pY-STAT1 and heterodimers with pY-STAT2 or pY-STAT3) to inhibit signalling via mislocalization and inhibition of DNA binding (21–24). In the effector stage, RABV-P protein can interact with the IFN-induced promyelocytic leukemia (PML) protein, impacting on PML nuclear bodies that are associated with the antiviral effects of IFN (25).

Many viruses, including members of the order *Mononegavirales* such as RABV, express proteins that can bind to and antagonise STATs, but the mechanisms of antagonism can differ, including inhibition of STAT phosphorylation (e.g. RABV), induction of degradation of STATs (e.g. Dengue virus), and arrest of STATs into high molecular complexes (e.g. Huaiyangshan banyangvirus) (14). Notably, RABV-P shows specificity in binding strongly to pY-STATs rather than unphosphorylated (U-) STATs (IFN/STAT-antagonists of many other viruses do not show such specificity), and does not induce degradation or inhibit phosphorylation, but rather prevents nuclear accumulation and DNA-binding (22–24, 26, 27). Inhibition of nuclear accumulation involves strong nuclear export of RABV-P-STAT complexes via a nuclear export sequence (NES) in RABV-P protein, while the inhibition of DNA binding appears to result from binding of RABV-P protein proximal to the DNA-binding site in STAT1 (23, 26, 28, 29). The RABV-P-STAT interface is conserved across members of the lyssavirus genus, which includes RABV and a number of other lethal viruses, and is critical in pathogenesis, such that RABV deficient for RABV-P-STAT1 interaction/antagonism is attenuated *in vivo* (30, 31). Thus, the RABV-P-STAT1 interface represents a potentially valuable target for the development of novel antiviral approaches including live attenuated vaccines against rabies, which is invariably lethal in symptomatic cases and results in an estimated > 60,000 human deaths/year (32).

Despite the established antiviral roles of IFN-α/β genes and importance of viral antagonism of the IFN response in infection and disease, ISGs can have ambivalent roles in infection. While some ISGs are extensively characterised for expected antiviral roles (e.g., MX1, PKR, APOBEC1, IFITM proteins, TRIM proteins), others have been shown to be ‘pro-viral’, with expression required for efficient infection (7, 33). Such roles vary between viruses presumably reflecting idiosyncrasies of the viral lifecycle (34). In RABV infection, the expression of heat shock protein (HSP70) and N-myc downstream-regulated gene (NDRG1) (both of which are reported to be induced by IFNs (35, 36)) correlates with increased RABV-P protein synthesis and viral budding respectively (37, 38). Similarly, the ISG B7-H1 is upregulated in RABV-infected neurons, dependent on type-I IFN, and B7-H1 knockout mice are resistant to infection with reduced disease severity, indicating that IFN-induced B7-H1 is required for infection (39). Thus, it would appear that RABV and other viruses are unlikely to benefit from a general antagonism of ISG expression, due to inhibition of both pro and antiviral genes. How viruses navigate the need to inhibit the expression of some ISGs while supporting others is poorly understood.

Although the type-I IFN response is generally considered in terms of pY-STAT1/2/IRF9 (ISGF3) signalling to regulate ISGs, it is considerably more complex. Type I IFNs can activate multiple STAT family members (STAT1-4, STAT5a, STAT5b and STAT6) as well as other non-canonical pathways, including PI3K and MAPK pathways (2, 40). U-STATs also regulate gene expression, and IFN-induced accumulation of U-STATs mediates extended regulation for distinct subset of ISGs (41). Furthermore, IFN can repress rather than activate the expression of a large number of IRepGs (7). Thus, selective targeting of specific elements of IFN signalling, such as pY-STAT1 complexes by RABV-P, would appear likely to affect selective regulation, rather than causing broad inhibition of the expression of ISGs. Importantly, recent data indicate that RABV-P binds to pY-STAT3/1 heterodimers, but not pY-STAT3/3 homodimers, in cells activated by the interleukin 6 (IL-6) cytokine family member oncostatin M (OSM), and this appears to relate to differential impacts on specific OSM-stimulated genes (FOS and SOCS3) (21). Thus, discriminatory targeting of immune signalling complexes by RABV-P protein might produce precise modulation of immune responses, possibly to favour the suppression of antiviral genes while supporting the expression of pro-viral genes. Delineation of this possibility requires system-wide analysis of the specific effects of IFN antagonists on the global IRG transcriptome, including antiviral or proviral ISGs and IRepGs.

Here, we report the use of RNA-sequencing and analysis including differential gene expression, pathway enrichment, TF prediction, and incorporation of public ChIP-seq data, to demonstrate that RABV-P protein selectively modulates IRG expression via specific TF targeting. RABV-P protein expression significantly affected the expression of only a subset of IRGs (including antagonising ISGs with known antiviral function), while permitting or enhancing the expression of other ISGs, including pro-viral genes. TF analysis indicates that RABV-P protein antagonises the effects of IFN on STAT1/2-regulated genes, as expected, but not genes regulated by ATF2, ELK1 and ELK4 TFs of the MAPK signalling pathway. Notably, RABV-P not only antagonised subsets of ISGs but also inhibited the suppression of subsets of IRepGs, highlighting the likely significance of such genes in the IFN-mediated antiviral response, and their modulation by viruses, an area that has remained poorly investigated.

## Result and Discussion

### RABV-P protein selectively regulates the expression of ISGs

To analyse the effects of RABV-P protein expression on the IFN-regulated transcriptome, we established stable integrated tetracycline (Tet)-inducible Flp-In™ T-REx™-293 cell lines expressing either FLAG-tagged RABV-P protein or FLAG-alone (control); the resulting cell lines were denoted 293-rex-FL-RABV-P and 293-rex-FL cells. This enabled rapid expression of FLAG-RABV-P or FLAG across the cell population, before treatment with or without type-I IFN (IFN-α) and analysis by RNA-seq (Figure 1). To validate and optimise the system, we assessed FLAG-RABV-P protein expression following induction with or without Tet (1 µg/ml, 24 h and 48 h) by immunoblotting of cell lysates for FLAG. FLAG-RABV-P protein expression was readily detectable in lysates of Tet-induced cells at both time points, with clearly increased expression compared with non-Tet treated cells (Supplementary Figure 1A). To confirm efficient expression across individual cells, we treated cells with Tet for 24 h before fixation, immunostaining for FLAG, and imaging by fluorescence microscopy. Results confirmed that Tet treatment induced broad expression of FLAG-RABV-P in the cells; furthermore, the localization of the anti-FLAG staining was consistent with expected largely cytoplasmic localization of RABV-P protein, due to the presence of a strong NES in RABV-P (Supplementary Figure 1B) (28). We further confirmed responsiveness of the cells to IFN by treatment of Tet-induced 293-rex-FL-RABV-P and 293-rex-FL cells with IFN-α for 1 h, 6 h or 24 h), followed by lysis and immunoblot (IB) analysis for pY-STAT1. pY-STAT1 was low or undetectable in 293-rex-FL-RABV-P and 293-rex-FL cells without IFN-α treatment and was rapidly induced following 1 h IFN-α treatment, consistent with the expected responses of cells to IFN-α and the observation that RABV-P protein does not impair STAT1 phosphorylation (Supplementary Figure 1C) (22, 42). Notably, while pY-STAT1 was lost over time in 293-rex-FL cells, consistent with expected dephosphorylation (42), dephosphorylation was inhibited in 293-rex-FL-RABV-P cells, such that pY-STAT1 levels were not reduced by 24 h post-IFN treatment. This is expected as RABV-P protein binding to pY-STAT1 inhibits dephosphorylation, as part of its IFN-antagonistic function (22, 26, 31, 42); thus, the data indicate that FLAG-RABV-P protein is functional and induced to expression levels sufficient to efficiently bind and antagonize cellular pY-STAT1. The lack of any apparent reduction in pY-STAT1 at later timepoints is consistent with efficient expression of FLAG-RABV-P protein across the cell population and lack of any significant sub-population lacking RABV-P cell protein expression as indicated by the microscopy data (Supplementary Figure 1B). To further confirm IFN-antagonistic function of Tet-induced FLAG-RABV-P protein, we used a well-established ISRE-dependent luciferase reporter gene assay to quantify type-I IFN-STAT1/2 signalling and effects thereon of RABV-P protein, as previously described (30, 42). Consistent with the results for IB of pY-STAT1 (Supplementary Figure 1C), IFN-α treatment of cells following Tet induction produced a clear increase in ISRE-dependent luciferase activity in 293-rex-FL cells, but luciferase activity in IFN-treated 293-rex-FL-RABV-P cells remained similar to untreated cells, indicating potent IFN/STAT1-antagonistic function (Supplementary Figure 1D).

**Figure 1:**
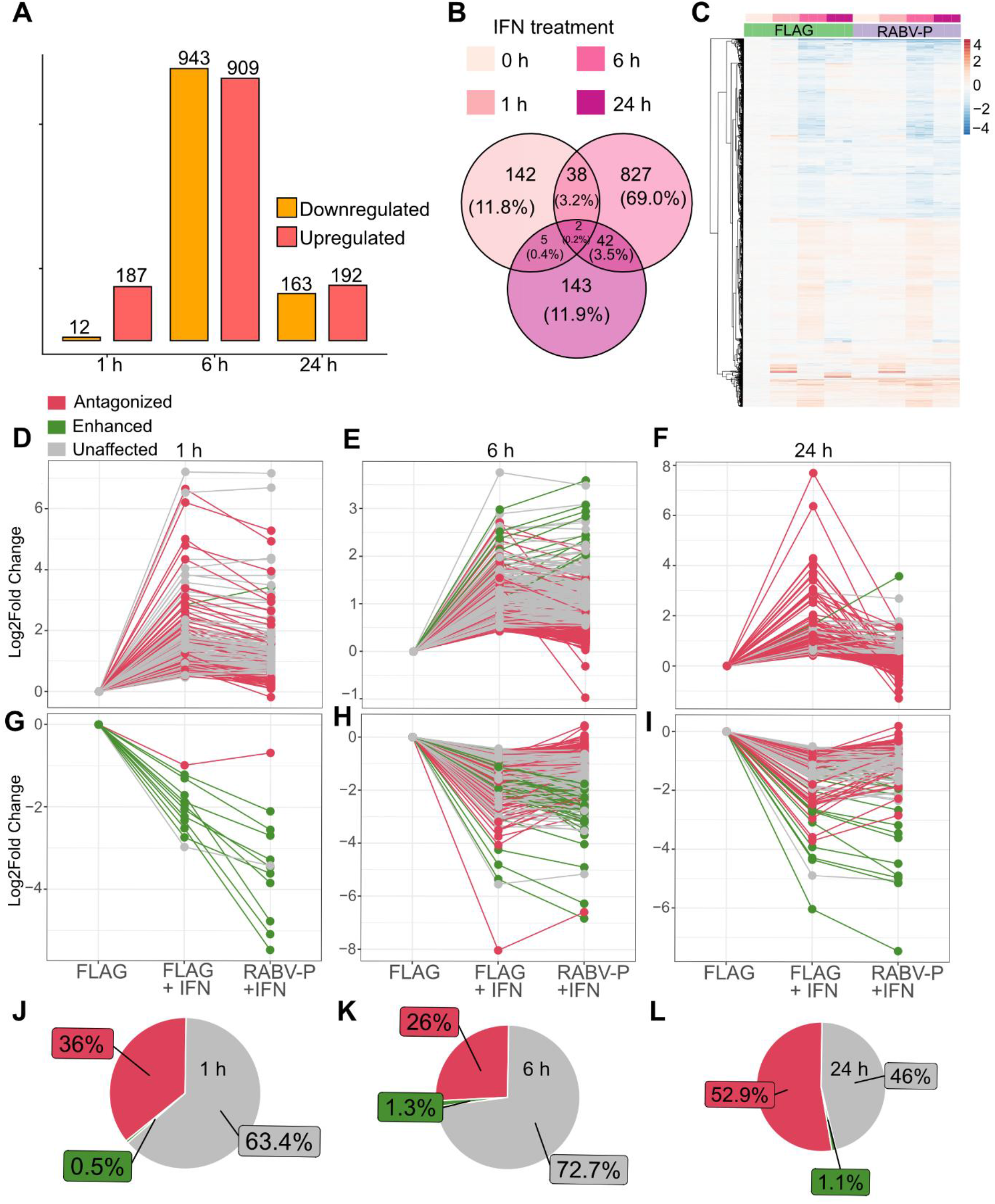
RABV-P protein expression inhibits a subset of ISGs: (A) Bar plots showing the number of genes significantly upregulated (red) or downregulated (orange) at 1 h, 6 h and 24 h post-IFN-α treatment in Tet-induced 293-rex-FL cells. (B) Venn diagram showing the overlap between significantly upregulated genes (treat threshold ≥1.2-fold change, FDR ≤ 0.05) in Tet-induced 293-rex-FL cells at 1 h, 6 h, and 24 h post-IFN-α treatment. (C) Heatmap of the average log₂ expression of genes in Tet-induced 293-rex-FL and 293-rex-FL-RABV-P cells at 1 h, 6 h and 24 h post-IFN-α treatment compared to 293-rex-FL cells without IFN-α treatment (0 h). Gene expression in all samples is normalized to the average levels observed in 293-rex-FL cells without IFN-α treatment. (D-I) Parallel coordinate plots of the log_2_ fold-change for ISGs (D-F) and IRepGs (G-I) at 1 h, 6 h, or 24 h post-IFN treatment showing whether the expression of each genes in Tet-induced 293-rex-FL-RABV-P cells (RABV-P +IFN) was antagonized (effect of IFN inhibited: gene expression not significantly induced or supressed exhibiting a decrease in IFN-induced log₂ fold change of > 0.5 compared with IFN-α treated 293-rex-FL cells (FLAG +IFN)); enhanced (effect of IFN increased: increase in log₂ fold change of >0.5 compared to IFN-treated 293-rex-FL cells), or unaffected (remained significantly induced, log₂ fold change of <0.5 compared with IFN-treated 293-rex-FL cells). (J-L) Proportions of significantly upregulated ISGs (FDR ≤ 0.05) that were antagonized, enhanced, or unaffected by RABV-P expression were determined by comparing the log₂ fold-change of ISGs in 293-rex-FL cells to those in 293-rex-FL-RABV-P cells, at 1 h (J), 6 h (K), and 24 h (L) post-IFN-α treatment.

We next collected and processed samples of 293-rex-FL cells following Tet induction (24 h) and treatment with IFN-α (0 h, 1 h, 6 h, 24 h) for RNA-seq analysis. RNA-seq data for Tet-induced FLAG-expressing cells were used to establish the baseline cellular response to IFN-α treatment in this system, by identifying ISGs and IRepGs across different timepoints, relative to untreated cells. Genes were considered significantly affected by IFN-α if they exhibited exhibiting a minimum 1.2-fold change (FC) and a false discovery rate (FDR) ≤ 0.05. This identified 187, 909 and 192 ISGs at 1 h, 6 h and 24 h timepoints post-IFN treatment (Figure 1A) (Supplementary Table 1). The ISG populations were largely distinct but with modest overlap between time points: 1 h to 6 h (38 shared genes), 6 h to 24 h (42 shared genes), and 1 h to 24 h (5 shared genes) (Figure 1B). Comparison with the Interferome database (36) revealed that the majority of ISGs identified had been previously characterised, including 65.8% (127/187) at 1 h, 45.5% (414/909) at 6 h, and 65.1% (125/192) at 24 h (Supplementary Table 1). Thus, our cell system demonstrated the ability to detect a large array of ISGs encompassing a diversity of IFN signalling outcomes, permitting analysis of the effects thereon of RABV-P expression.

To assess the effect of RABV-P protein on the global expression of ISGs, we compared ISG expression between Tet-induced 293-rex-FL and 293-rex-FL-RABV-P cells, at 1 h, 6 h and 24 h post-IFN-α treatment (Figure 1C). The majority of ISGs exhibited similar expression patterns in 293-rex-FL-RABV-P and 293-rex-FL cells (Figure 1 C, D-F, J-L), indicating that RABV-P does not cause a global suppression of ISGs.

We next compared the expression of the subset of genes that were significantly upregulated (FDR ≤ 0.05) in 293-rex-FL cells with their expression in 293-rex-FL-RABV-P cells at each time point post-IFN treatment. This indicated that only 36% and 26% of ISGs were antagonized by RABV-P expression at 1 h and 6 h, respectively, with 63.4% and 72.7% of the remaining ISGs being unaffected by RABV-P expression (Figure 1 J, K). At 24 h post-IFN-α treatment, a higher proportion of ISGs were antagonized (52.9%) than unaffected (46%) by RABV-P (Figure 1 L). A small subset of ISGs also appeared to display enhanced expression in 293-rex-FL-RABV-P cells at 1 h (0.5%), 6 h (1.3%), and 24 h (1.1%) IFN-α treatment (Figure 1 D-F, J-L); however, the limited number of genes identified constrained statistical power to attribute this effect to RABV-P protein expression.

As the ISGs at different timepoints were largely distinct, the observation that RABV-P impacts different proportions of ISGs at different timepoints may relate to the involvement of different cellular gene regulatory pathways over the course of the experiment. Specifically, regulatory pathways at the earliest timepoint (1 h), which is likely to include “early” gene expression that occurs rapidly in response to stimulation, without the requirement for new protein synthesis (43), is more sensitive to RABV-P protein than pathways affecting gene expression at 6 h post-IFN-α treatment (see below). The observation of increased ISG antagonism at 24 h suggest an increase in regulatory pathways affected by RABV-P protein, which may be expected as many ISGs are components (e.g. STATs, IRFs) or regulators of classical IFN signalling, which form feedback loops (e.g. SOCS1/IRF7), some of which are known to be inhibited by RABV-P protein. Overall, these data indicated that RABV-P protein selectively suppresses the expression of a distinct sub-population of ISGs.

### Analysis of ISGs regulated by RABV-P protein

To understand the biological functions and pathways associated with the ISGs antagonized by RABV-P, we performed gene set enrichment analysis (GSEA). Analysis of ISGs induced at 1 h post-IFN-α treatment in 293-rex-FL cells revealed a significant enrichment for genes regulated by the NF-κB pathway (TNF-α signalling via NF-κB) (FDR = 1.4e-67, avg. log_2_FC = 0.88), which was inhibited in 293-rex-FL-RABV-P cells (FDR = 2.4e-24, avg. log_2_FC = 0.67) (Figure 2A), including several early response genes, such as the MAPK signal transduction mediators DUSP1 (log_2_FC = 2.57, 1.52 (293-rex-FL, 293-rex-FL-RABV-P respectively)), the NF-κB signalling regulator NFKBIZ (log_2_FC = 2.53, 1.89) ; AP-1 complex members FOS (log_2_FC = 5.01, 3.1) and FOSL2 (log_2_FC = 1.7, 0.88); interferon regulatory factor 1 (IRF1) (log_2_FC = 3.08, 1.7); members of the early growth response family EGR1 (log_2_FC = 4.8, 3.9), EGR2 (log_2_FC = 6.2, 5.2) and EGR4 (log_2_FC = 6.6, 4.9); and ZFP36 (log_2_FC = 3.4, 2.6), an RNA-binding protein that promotes the degradation of mRNAs containing AU-rich elements (44–46). These findings indicate that RABV-P protein modulates the expression of early response genes, including those also regulated through the NF-κB pathway, with several of these genes known to play antiviral roles.

**Figure 2:**
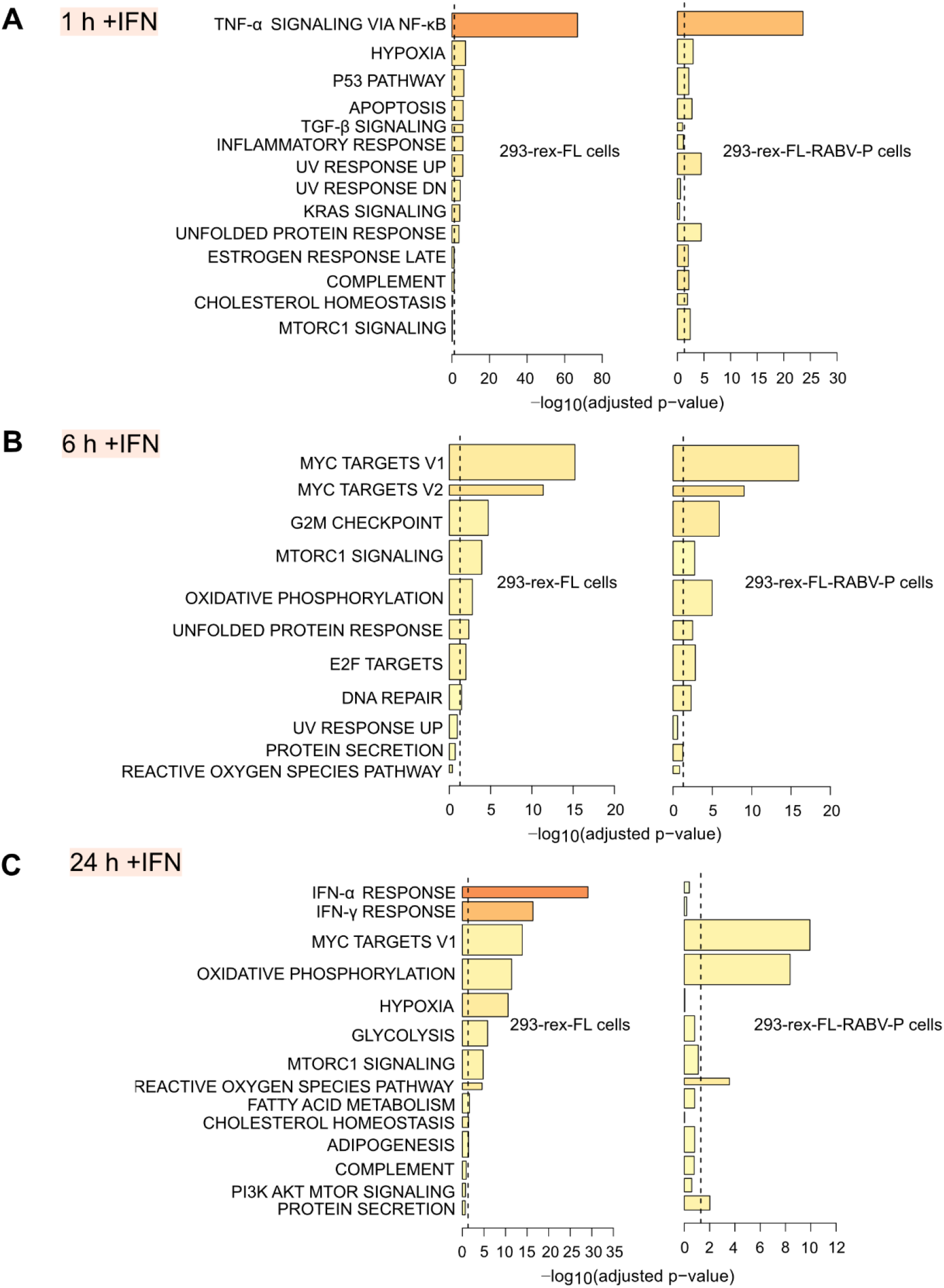
GSEA analysis of IFN-treated 293-rex-FL and 293-rex-FL-RABV-P cells: Bar plots representing the results of GSEA analysis for Tet-induced 293-rex-FL and 293-rex-FL-RABV-P cells at the indicated times post-IFN-α treatment, using the MSigDB hallmark gene sets as reference. Each panel compares pathway enrichment in cells expressing the control FLAG protein (left; 293-rex-FL) or RABV-P (right; 293-rex-FL-RABV-P). Bar length represents the -log_10_ adjusted *P* value, colour indicates the mean log_2_ fold change of genes within each gene set (red: upregulated; blue: downregulated), and bar width corresponds to the relative size of the gene set. The dashed line indicates the FDR significance threshold (adjusted *P* = 0.05). Results are ordered by significance in 293-rex-FL cells.

At 6 h post-IFN-α treatment, GSEA analysis of genes induced in 293-rex-FL cells and 293-rex-FL-RABV-P cells revealed broadly similar pathway enrichment profiles (Figure 2B), consistent with differential expression analysis showing that RABV-P does not affect the expression of the majority (72.7%) of ISGs at this timepoint (Figure 1 K). Notably, the subset of ISGs antagonized by RABV-P included well characterized antiviral genes such as IRF9 (log_2_FC = 1.9, log_2_FC = -0.3), IFI44L (log_2_FC = 2.7, log_2_FC = 0.4) (47, 48), and proteins associated with the secretory pathway like SCG3 (log_2_FC = 1.5, log_2_FC = 0.9) (Supplementary Table 1). Thus, RABV-P protein appears to impact on specific ISGs that include genes with antiviral roles, and genes with roles in other cellular processes with potential importance to infection (i.e. secretory pathway and enveloped virus assembly/budding) (47–49). RABV-P protein also supressed classical antiviral ISGs at the 24h timepoint, where GSEA indicated significant enrichment for components of the IFN-α and IFN-γ pathways in 293-rex-FL cells that was absent in 293-rex-FL-RABV-P cells (Figure 2 C). These genes included well-defined antiviral effectors IFIT1 (log_2_FC = 4.2, log_2_FC = 0.00), IFIT2 (log_2_FC = 3.6, log_2_FC = -0.5), IFIT3 (log_2_FC = 6.3, log_2_FC = -0.5), IFI6 (log_2_FC = 3.6, log_2_FC = - 0.99), IFI44L (log_2_FC = 7.6, log_2_FC = 1.5), ISG15 (log_2_FC = 3.9, log_2_FC = 0.11), and OAS3 (log_2_FC = 1.8, log_2_FC = 0.15). The list of genes inhibited by RABV-P also included genes involved in propagating/amplifying the IFN response through positive feedback pathways, such as IRF9 (log_2_FC =3.4, log_2_FC = 0.1) and STAT1 (log_2_FC = 2.5, log_2_FC = -0.2), and RNA processing and regulation factors RIG-I (log_2_FC = 2.9, log_2_FC = -0.2) and DDX60 (log_2_FC =2.9, log_2_FC = 0.93), which are involved in viral RNA sensing and activation of the IFN-induction pathway. Overall, the results indicated that RABV-P targets pathways that stimulate ISGs important to establishing and maintaining the antiviral state.

While these data support the importance of IFN-antagonistic function of RABV-P in inhibiting antiviral responses, a large proportion of ISGs were not affected by RABV-P (63.4%, 72.7%, 46% of ISGs at 1 h, 6 h, 24 h respectively, Figure 1J-L). This suggests that IFN regulation is mediated through multiple pathways with only a subset affected by RABV-P. Thus, it is plausible that this selective antagonism enables the virus to avoid supressing the expression of genes that do not have antiviral roles or may be required or beneficial (pro-viral) for viral infection. Consistent with this, we found that at 1 h post IFN-α treatment, several genes encoding TFs that negatively regulate the type-I IFN response, including KLF4 (log_2_FC = 1.9, log_2_FC = 1.7), MAFB (log_2_FC = 1.7, log_2_FC = 1.4), SPRYR (log_2_FC = 1.1, log_2_FC = 1.2), and ZC3H12A (log_2_FC = 3.5, log_2_FC = 3.2) (50–53) were upregulated in 293-rex-FL cells and not inhibited in 293-rex-FL-RABV-P cells. The sustained expression of these negative regulators in 293-rex-FL-RABV-P cells may represent a mechanism by which the virus exploits host feedback pathways to dampen the antiviral state. Other ISGs whose expression was not inhibited by RABV-P protein included several genes implicated in apoptosis (e.g. GADD45G, MCL1, CASP3) (Supplementary Table 1). The regulation of apoptosis has been implicated in RABV infection with several studies demonstrating that RABV induces apoptosis in the host cell involving various mechanisms; these data suggest a possible contribution by certain ISGs (54–56). Together these findings suggest that RABV-P does not inhibit the expression of a number of genes involved in processes such as negative regulation of the type-I IFN response and apoptosis, indicating that these pathways may remain functionally active and contribute to a proviral environment.

Both 293-rex-FL and 293-rex-FL-RABV-P cells showed enrichment for genes involved in oxidative phosphorylation and unfolded protein response (UPR) pathways at 6 h and 24 h post-IFN-α treatment (Figure 2B, C). This suggests that RABV-P expression does not significantly impact on the effects of IFN on cellular energy metabolism and UPR signalling. Consistent with the enrichment of proteins involved in protein folding, several heat shock proteins (HSPs) involved in chaperone-mediated protein folding, including HSPA8 (log_2_FC = 2.24, 2.42 (293-rex-FL, 293-rex-FL-RABV-P respectively)), HSPA1A (log_2_FC = 3.76, 3.49), HSPA5 (log_2_FC = 1.74, 1.48), DNAJB1 (log_2_FC = 1.93, 2.07), and HSPH1 (log_2_FC = 1.70, 1.73), were upregulated by IFN to similar extents in 293-rex-FL and 293-rex-FL-RABV-P cells. HSPs are induced in response to several cellular stressors, including viral infections and facilitate the activation of immune signalling pathways. However, they are also hijacked by several viruses to aid replication and maturation (57). Notably, the expression of genes encoding members of the HSP70 and HSP90 families (HSPA8 and HSP90B1, respectively) was not affected by RABV-P (Supplementary Table 1). Previous studies have reported that HSP70 and HSP90 positively regulate RABV infection (37, 58), providing examples that are consistent with a model in which selective modulation of gene expression by RABV-P avoids supressing proviral host factors in IFN-activated cells.

Other ISGs expressed in 293-rex-FL and 293-rex-FL-RABV-P cells and not significantly affected by the expression of RABV-P protein include, NDRG1 (log_2_FC = 1.5, 1.3 (293-rex-FL, 293-rex-FL-RABV-P respectively)), which facilitates RABV infection by enhancing viral budding, and DYNLL1/LC8 (log_2_FC = 1.07, 2.11), a dynein-light-chain subunit that binds a dynein-light chain-association sequence in RABV-P to promote viral transcription and replication, and is involved in nuclear impirt and nucleolar localization of RABV-P (38, 59–64). Additionally, we identified several ISGs that were not inhibited by RABV-P and have been reported to facilitate infection by other viruses, such as KLF4 (log_2_FC =1.9, 1.7), which induces differentiation dependant lytic Epstein-Barr virus infection (65) and PCDH10 (log_2_FC = 2, 3.1), the neuronal receptor for Western equine encephalitis virus (66); whether these have roles in RABV infection is unclear, but are consistent with the existence of a broad range of pro-viral ISGs.

These data collectively indicate that the RABV-P protein modulates the IFN signalling system to inhibit certain ISGs while not affecting the expression of a diverse array of distinct ISGs. The findings further suggest that similar selective antagonism of IFN signalling pathways by other viruses may produce a similar balance of effects on anti- and pro-viral genes, including regulators of biosynthesis, immunity and cell survival with broad cellular effects impacting on the obligate intracellular replication of viruses, or with more direct roles in supporting replication processes. Overall, these data support the idea that, while proteins such as RABV-P protein are referred to as IFN antagonists (14, 16, 67), these proteins (including those that inhibit ‘core’ mediators such as STAT1) can be highly selective in their effects on IFN signalling, specifically modulating rather than globally inhibiting responses.

### Selective targeting of transcription factors by RABV-P protein correlates with differential regulation of ISGs

The above data indicate that RABV-P selectively impacts on pathways that regulate ISGs. The observation that largely distinct sets of ISGs are activated by IFN-α at the different time points (Figure 1B), indicates that multiple regulatory pathways and TFs contribute to ISG regulation across the time course, with the capacity of RABV-P to antagonise distinct ISG subsets (Figure 1D-L) suggestive of potential antagonism of several pathways. To assess the mechanisms underlying selective ISG antagonism we initially sought to identify the TFs associated with ISG expression at each time point, using an enrichment-based TF binding site (TFBS) analysis. Specifically, we analysed the promoter regions in those ISGs significantly upregulated at each timepoint (FDR ≤ 0.05; 1 h, 6 h, 24 h) in 293-rex-FL cells, to identify TFBS enriched relative to those in genes that were expressed but not significantly upregulated.

This analysis identified multiple TFs with largely distinct sets identified for each time point, reflecting the complex and dynamic nature of IFN-dependent signalling (Figure 3A-F, Supplementary Table 2). Notably, only a limited number of TF families were shared across timepoints (e.g., the Kruppel-like factor (KLF) family at 1 h and 24 h); instead, most appeared uniquely at specific time points, (e.g., the ETS family at 6 h, and STAT family at 24 h), indicating time-dependent ISG regulation by different TF families during IFN-α responses.

**Figure 3:**
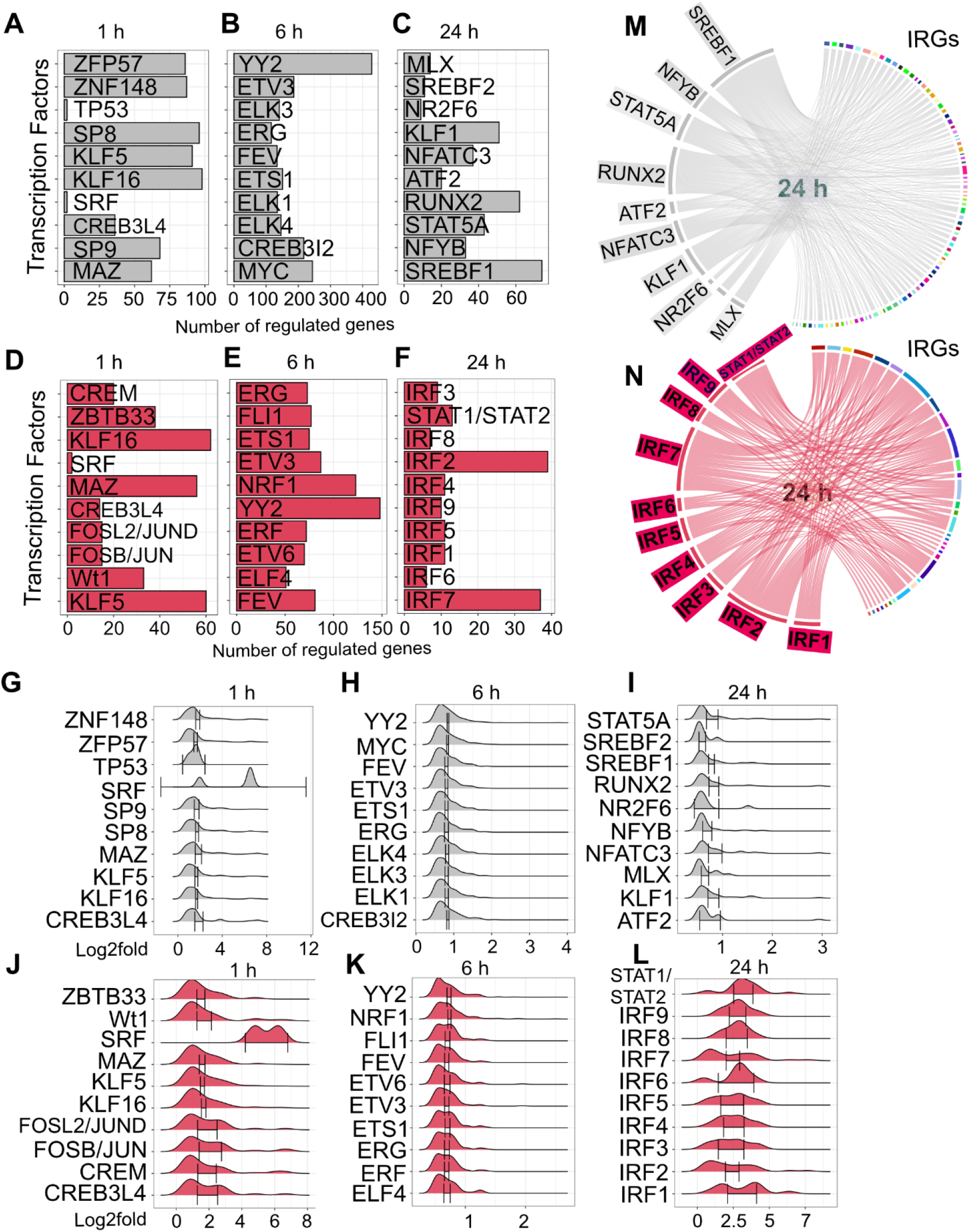
Predicted TFs regulating ISGs affected or unaffected by RABV-P expression: CiiiDER enrichment analysis of ISGs either antagonized (red) or unaffected by RABV-P (grey). (A-E) Bar plots showing the top ten TFs ranked by significance score, indicating their likelihood of being true regulators with the number of target genes at 1 h, 6 h, and 24 h post-IFN-α treatment. (G-L) Ridge line plots showing the distribution of expression changes for the target genes of each predicted TF. The x-axis shows the log2 fold change in gene expression, whereas the y-axis lists TFs. Density curves represent the distribution of log2 fold changes across the target genes regulated by each TF. Right-shifted curves indicate stronger overall ISG induction, whereas broader curves reflect greater heterogeneity in the ISGs regulated. (M, N) Representative Circos plots depicting the regulatory networks between predicted TFs and ISGs at 24 h post-IFN-α treatment. The left side of each ring shows TFs, and the right-side shows ISGs, with ribbons indicating predicted regulatory relationships. Ribbon thickness corresponds to the number of inferred interactions.

Having identified potential TFs at each time point, we sought to independently validate binding of these TFs to the promoter regions of their respective ISG targets. For this we used the public ChIP-seq database ENCODE (68) to assess the strength of TF binding at specific genomic locations. This confirmed binding for 41 of the predicted 51 TFs at their ISG targets (Supplementary Table 3). Our subsequent analysis prioritized these confirmed TFs.

RABV-P protein binds to directly to pY-STAT1, and inhibits transcription by pY-STAT1 homodimers, and pY-STAT1/2 and pY-STAT1/3 heterodimers in response to IFNs and other cytokines (21, 23, 26, 69). Consistent with this, our results showed an enrichment for STAT1/2 binding sites in genes antagonized by RABV-P at the 24 h timepoint (Figure 3F, L, N). Our analysis also identified multiple members of IRF family (IRF1-IRF9) as potentially regulating the expression of genes antagonized by RABV-P (Figure 3F, L, N). Several IRFs have been reported to be activated by IFN (70) and RABV-P protein inhibits the activation of IRF3 and IRF7 in the IFN induction pathway (71, 72); thus, our data suggest that inhibition of IRF3/7 may contribute to antagonism of IFN signalling as well as IFN induction. The finding that RABV-P protein inhibits genes that are largely regulated by STATs/IRFs at the 24h time point is consistent with the idea that RABV-P antagonises a specific subset of IRGs via a highly selective interaction with IFN-activated pY-STAT1 and suggests that inhibition of specific IRFs also contributes.

Activation by IFN of non-canonical pathways (e.g. MAPK, PI3K, MTOR and STAT1/2 independent pathways), in addition to the classical STAT1/STAT2 dependent pathway, may account for the differential regulation of ISGs by RABV-P; however, the influence of RABV-P protein on these pathways remains largely unexplored. Our analysis of ISGs inhibited at the 6 h timepoint indicates that a subset of TFs including ELF4, FLI1, ETS1, ETV3 and ETV6 were enriched in promoter sites of ISGs inhibited by RABV-P (Figure 3E, K). Among these, ELF4 is a well-characterized antiviral transcription factor that plays a critical role in type-I IFN induction and host antiviral defence (73). ELF4 is phosphorylated by TBK1, enabling its nuclear translocation and binding to ISG promoters, where it cooperates with IRF3 and IRF7 to drive transcription (73). Knockdown of ELF4 reduces IFN production and impairs IRF3/IRF7 binding at ISG promoters, leading to increased susceptibility to virus infection. Given that RABV-P inhibits TBK1-dependent phosphorylation of IRF3/7 (20), it is plausible that RABV-P may also inhibit activation of ELF4 via a similar mechanism. Similarly, members of the ETS TF family, ETS1 and FLI1, have been implicated as key regulators of IFN responses. FLI1 is activated downstream of IFN signalling and promotes STAT1 expression in response to IFN-γ (74), while ETS1 is essential to immunity through its regulation of IL-12 receptor expression and promotion of IFN-γ production (75, 76). In contrast, ETV3 and ETV6, which function primarily as transcriptional repressors within cytokine signalling networks, may act to fine-tune the amplitude of ISG expression (77). Thus, the direct or indirect inhibition of these TFs by RABV-P could interfere with both the activation and repression of IRGs, thereby enabling the virus to dampen the antiviral state by modulating multiple pathways.

TFBS enrichment analysis on the promoter sites of the ISGs antagonized by RABV-P at 1 h IFN treatment, identified enrichment of binding motifs for CREM, ZBTB33, KLF16, SRF, MAZ, CREB3L4, FOSL2-JUND, FOSB-JUN, WT1, and KLF5 (Figure 3D, J). Many of these TFs are key regulators of immediate-early gene expression and cellular stress responses (78), suggesting that RABV-P may interfere with early transcriptional events downstream of IFN signalling. For example, MAZ controls the expression of antiviral genes such as IRF8, IFIT2, and ISG15 in response to IFN-γ treatment and is essential to the ability of STAT1’s to bind to its genomic targets (79). Another early response TF, SRF, regulates the expression of ISGs downstream of type-I IFN signalling but operates independently of the classical STAT1/2 regulated pathway (80). The enrichment of KLF5 and KLF16, members of the Krüppel-like factor family, implicates RABV-P in modulating PI3K/mTOR and cell-cycle regulatory networks that act downstream of type-I IFN signalling (2, 81). Collectively, these findings suggest that RABV-P antagonism extends beyond inhibition of STAT1 to target a broader network of MAZ/KLF/SRF-driven transcriptional programs, thereby dampening a number of pathways involved in the immediate-early phase of IFN-induced gene activation.

TFBS analysis of ISGs unaffected by RABV-P identified several TFs distinct from the TFs identified for ISGs antagonized by RABV-P. This was particularly evident at 24 h post-IFN treatment, where all TFs associated with unaffected ISGs (Figure 3C, I, M) were unique from those suppressed by RABV-P (Figure 3F, L, N). Among the unaffected TFs at 24 h post-IFN-α treatment, MLX regulates lipid and glucose metabolism (82), while SREBF1 and SREBF2 regulate lipid biosynthesis (83, 84), which suggests that the maintenance of activity of these TFs may support RABV infection. SREBP2 activated gene expression has been shown to enhance Zika virus (ZIKV) infection (85). Although the pathogenic mechanism of ZIKV and RABV differ substantially, these findings suggest that SREBF mediated gene regulation may also influence RABV infection, warranting further investigation. We also identified transcriptional repressors such as NR2F6 (which acts as an immune checkpoint–like regulator to antagonize the expression of cytokines including IL-2 and IFN-γ (86)) and ATF2 (which represses the expression of IFN-β and downstream type-I IFN signalling (87)). We also identified TFs that are activated by and contribute to antiviral IFN response such as NFATC3 and STAT5A (14, 88, 89). The role of STAT5A in viral infection appears complex, exhibiting both proviral and antiviral activities depending on the virus (14), however these data suggest it may have proviral effects for RABV. Similarly, NFATC3 regulates the cell cycle and apoptosis can induce ISG expression (89), but is also required for infection by the DNA virus human polyomavirus JC virus (90), underscoring its pleiotropic functions, which may vary between different viruses. Together these findings suggest that TFs that are not targeted by RABV-P include TFs with roles in metabolism and repression of IFN-mediated immune responses.

We identified enrichment of TFBS for oncogenic TFs, including the ETS family (ELK1, ELK3, ELK4, ETS1) and MYC, in the promoter sites of ISGs unaffected by RABV-P at 6 h post-IFN treatment (Figure 3B, H). These TFs are key regulators of cell growth, proliferation and survival, and are exploited by certain viruses to enhance viral gene expression and sustain host cell viability (91, 92). Thus, it is plausible that the preservation of the activity of these TFs may also support cellular homeostasis and preserve cellular integrity to provide a favourable environment for RABV infection. This aligns with the characteristic pathology of pathogenic RABV infection, which causes minimal cytopathic effects in infected neurons and limited tissue damage (93, 94).

Similarly, enrichment of TFBS for ZFP57, ZNF148, TP53, SP8, and SP9 among ISGs unaffected by RABV-P at 1 h post-IFN treatment (Figure 3A, G), suggests that RABV-P avoids disruption of TFs essential for neuronal function and survival. SP8 and SP9 regulate the development, migration, and differentiation of interneurons (95–97), so maintenance of their function may be linked to maintaining the integrity of neurons during infection. TP53 regulates cell fate by controlling cell cycle arrest, apoptosis, and DNA repair (98) and regulates the type-I IFN response playing a key role in antiviral innate immunity (99, 100). Despite reports of antiviral roles, certain viruses activate or upregulate p53 (100), implying that its activity can be co-opted to serve virus-specific processes. Together, these observations suggest that TFs not inhibited by RABV-P include TFs with roles in cell homeostasis, including neuronal transcriptional programs, suggesting that RABV-P preserves transcriptional programs important for cell survival and neuronal function.

### RABV-P protein can reverse the suppression of specific IRepGs

The IFN-dependent antiviral system is conventionally considered in terms of the activation of antiviral ISGs, with viral IFN antagonist studies focusing on the inhibition of ISG induction (7, 10). However, IFN represses a substantial number of IRepGs (7). We confirmed significant suppression (FDR ≤ 0.05) of subsets of IRGs (12, 943, and 163 IRepGs at 1 h, 6 h, and 24 h post-IFN treatment, respectively) in 293-rex-FL cells (Figure 1A, G, H, I) (Supplementary Table 1). Notably the number of IRepGs at 6 h moderately exceeded the number of ISGs, consistent with previous findings (11). Importantly, our data indicated that RABV-P protein antagonized IFN-mediated repression at all investigated timepoints, affecting 8.3% (1/12), 35.1% (331/943), and 35.6% (58/163) of IRepGs at 1 h, 6 h, and 24 h, respectively. This suggests that RABV-P interferes with regulatory mechanism governing both ISG activation and IRepG repression, which may involve the same regulatory pathways that have distinct (stimulatory or repressive) effect on different genes. Analysis using the Interferome database confirmed that a substantial proportion of IRepGs identified in our RNA-seq data correspond to previously reported IRGs, including 33.3% (4/12), 64.7% (610/943) and 52.2% (90/163) of IRepGs at 1 h, 6 h, and 24 h, respectively. Collectively, these data indicate that RABV-P selectively antagonizes IFN-mediated gene repression, as well as ISG activation.

While less well understood than the induction of ISGs, the suppression of IRepGs by IFN appears to contribute to the host antiviral response by limiting host dependent factors (e.g. FASN, Ergic53, IDH1) known to be important to infection by viruses including HIV, dengue virus and hepatitis C virus (11). Although our literature review did not identify any of the IRepGs from the current study as being directly implicated in RABV infection, several possess pro-viral functions that could potentially facilitate viral infection if their expression is restored by the IFN antagonistic function of RABV-P. For example, WNT3A, a key ligand in the canonical Wnt/β-catenin signalling pathway, regulates cellular communication, development, and immune modulation, and has been shown to be enriched in astrocytes that support high levels of RABV mRNA transcription (101).

Given the importance of STAT1 and STAT2 TFs in type-I IFN signalling and their antagonism by RABV-P to inhibit ISG activation, it is possible that this antagonism extends to potential gene repressor activities of STAT1/2. However, in our analysis, most IRepGs lacked canonical STAT1 or STAT1/2 binding motifs, suggesting that these sites may be primarily associated with transcriptional activation rather than repression (9). These results are similar to a study on IFN-γ-induced repression of genes which did not find any significant enrichment for STAT1/2 (ISRE), STAT1/1 (GAS) or IRF sites in the promoter region of IRepGs, despite the fact that repression was lost in STAT1-deficient cells compared to wild type cells (11). This implies that STAT1 mediates gene repression through motif-independent mechanisms, possibly through the induction of regulators. Additional mechanisms are also likely contribute to IFN-induced gene repression, including epigenetic modifications, inactivation of enhancers and post-transcriptional regulatory pathways (102, 103). Future studies will be required to elucidate the mechanisms underlying IRepG suppression by RABV-P and other IFN antagonists, including the contribution to antagonism of targeting specific TFs such as STAT1 and newly predicted RABV-P-regulated TFs indicated in this study. Nevertheless, the observed enrichment of STAT1 sites in ISGs antagonized by RABV-P and well established interface of RABV-P with STAT1(24, 27), together with the known involvement of STAT1 in IRepG repression (9) and the similar trend of antagonism of ISGs and IRepGs at different timepoints, suggest that the effect of RABV-P on ISGs and IRepGs may involve targeting of the same signalling pathways, particularly those involving pY-STAT1.

In conclusion, our data indicate that RABV-P effects only partial antagonism of IRG expression, selectively inhibiting effects of IFN on a subset of IRGs including antagonising classical antiviral ISGs, while permitting the expression of other subsets of ISGs, including ISGs with known proviral roles for RABV. Consequently, it appears that IFN antagonists can generate a coordinated effect not only suppressing a hostile cellular environment but also fostering a cellular environment supportive of infection.

## Material and Methods

### Cell lines

Cell lines for Tetracycline (Tet) inducible expression of RABV-P protein were generated using the Flp-In T-REx™-293 system (Invitrogen) according to the manufacturer instructions. Briefly pcDNA 5/FRT/TO (Invitrogen) constructs containing an insert to express FLAG or FLAG-tagged RABV-P (generated by PCR amplification from a plasmid containing the RABV-P of the challenge virus standard (CVS) strain of RABV(30)) were co-transfected with pOG44 plasmid which carries a flippase enzyme for integration of the inserted genes into the genome. Transfection was performed using Lipofectamine 2000 (Invitrogen), according to the manufacturer’s instructions. Post-transfection, cells were maintained in Dulbecco’s modified Eagle’s medium (DMEM; Gibco) supplemented with 10% fetal bovine serum (Gibco), blasticidin (25 μg/ml; Sigma-Aldrich), and hygromycin B (100 μg/ml; Sigma-Aldrich) at 37 °C with 5% CO_2_ in a humidified incubator. The integrated cell lines were designated 293-rex-FL and 293-rex-FL-RABV-P.

### Analysis of protein induction

To assess Tet induction of FLAG-RABV-P expression, 293-rex-FL and 293-rex-FL-RABV-P cells were treated with or without Tet (1 μg/ml) for 24 h and 48 h. For Immunoblot (IB) analysis, cells were washed with PBS and harvested and lysed in lysis buffer (10 Tris, 150mM NaCl, 0.5 mM EDTA, 0.5% NP40), supplemented with complete protease inhibitor and PhosSTOP^™^ (Sigma-Aldrich). IB analysis used anti-FLAG (Sigma-Aldrich, F1804), anti-β tubulin antibody (Sigma-Aldrich, T8328) and goat anti-mouse HRP-conjugated secondary antibody, with detection using ECL lightening Plus reagent (Revvity).

For analysis of FLAG-RABV-P expression by microscopy, cells treated for 24 h with Tet (1 μg/ml) were fixed with 4% formaldehyde in PBS (15 min, room temperature) before permeabilization with 0.5 % Triton X-100 in PBS (5 min, room temperature) and blocking using 10% bovine serum albumin in PBS (60 min, room temperature). Cells were then immunostained using anti-FLAG antibody (Sigma-Aldrich, F1804) followed by anti-mouse Alexa Fluor 568 (Thermo Fisher) and imaged using an EVOS Cell Imaging System (Invitrogen); nuclei (DNA) were visualised using Hoechst 33342.

### Analysis of IFN signalling and IFN antagonist function of FLAG-RABV-P protein

To confirm the activity of IFN-α-dependent STAT1/2 signalling in 293-rex-FL and 293-rex-FL-RABV-P cells, and antagonistic function of Tet-induced FLAG-RABV-P, we used IB (above) and an IFN-α STAT1/2 dependent dual luciferase reporter gene assay (30, 42). For IB, 293-rex-FL and 293-rex-FL-RABV-P cells were induced with Tet (1 μg/ml, 24 h) before treatment with or without IFN-α (1000 U/ml) for 1 h, 6 h,16 h, or 24 h. Cells were lysed and analysed by IB (as above) using anti-FLAG (Sigma-Aldrich, F1804), anti-tubulin (Sigma-Aldrich, T8328) and rabbit anti-pY-STAT1 (CST #9167) antibodies, followed by HRP-conjugated anti-rabbit secondary antibodies and detection with the ECL lightening Plus reagent (Revvity).

For luciferase reporter gene assays, 293-rex-FL and 293-rex-FL-RABV-P cells were transfected with plasmids for the IFN-α STAT1/2 dependent dual luciferase reporter gene assay (pRL-TK) (40 ng), which expresses *Renilla* luciferase constitutively and pISRE-luc (250 ng), which expresses firefly luciferase under the control of an ISRE-containing promoter) before induction with Tet (1 μg/ml, 24 h). Cells were then treated with or without IFN-α (1000 U/ml, 16 h). Cells were then lysed with a passive lysis buffer (Promega) and analysis of lysates by a dual luciferase assay, as previously described (28, 30, 42). Relative luciferase activity (RLU) was calculated by dividing the firefly luciferase values by the *Renilla* luciferase values.

### RNA extraction and sequencing

RNA was extracted from Tet-treated (1 μg/ml, 24 h) 293-rex-FL and 293-rex-FL-RABV-P cells following with or without IFN-α (1000 U/ml, 1 h, 6 h, 24 h), using an RNeasy RNA extraction kit (Qiagen); this included an on-column DNAase digestion using the RNase-free DNAse kit (Qiagen), according to the manufacturer’s instructions. Following extraction, the RNA integrity number (RIN) was determined using a Bioanalyzer (Agilent Technologies) before proceeding to RNA sequencing.

Libraries were generated by in-house multiplex RNAseq method (Hudson Genomics Facility) using 20 ng of total RNA input. The library is designed so that a custom forward read (R1 19 bp) primer sequences the sample index, followed by a 10bp unique molecular identifier (UMI). The reverse read (R2 72bp) primer sequences the cDNA in the sense direction for transcript identification. Sequencing was performed on a NextSeq550 (Illumina) platform using the v2.5 High Output Kit. Base calling was performed using Bcl2fastq (v2.20.0.422, Illumina).

### Bioinformatic analysis

RNA-seq analysis was performed in R (v4.2.2). Sequencing reads were first processed and demultiplexed using the R package ScPipe (v1.16.0) (104). The sc_trim_barcode function was used to reformat and combine R1 and R2 FASTQ files by trimming the sample index and UMI sequences and storing them as the read header (with bs2 = 0, bl2 = 8, us = 8, ul = 10). Reads were aligned to the human primary assembly genome file using the Rsubread (v3.17) package (105).

Following alignment reads were mapped and annotated by using the Ensembl Homo sapiens GRCh38 v111 GFF3 genome annotation file through the sc_exon_mapping function. The resulting BAM file was demultiplexed using the sc_demultiplex function, which associates reads mapped to exons with each individual sample. Subsequently, the sc_gene_counting function generated an overall gene count for each sample.

Gene counts were further annotated using the biomaRt (v2.58.2) package. A DGElist consisting of the gene counts and their annotation was then created with the edgeR package (v4.0.14). Before further analysis, lowly expressed genes were filtered out using the filterByExpr function and normalisation factors were calculated using the TMM method (106). Samples were arranged into groups, and gene counts were transformed by modelling a mean-variance relationship and applying precision model weights to each gene using the voom function. A linear model was fitted using the lmFIt function from the limma package (v3.58.1) (107). The contrasts.Fit function was used to compare groups, and moderated t statistics were calculated using the treat function at a 1.2-fold change threshold with differentially expressed genes determined using an adjusted false discovery rate of *P* < 0.05. Heatmaps were created using the pheatmap package (v1.0.12). Relative Log_2_CPM values were calculated by subtracting the mean log_2_CPM value of the untreated FLAG control samples from the corresponding log_2_CPM values for each gene.

Gene set enrichment analysis was performed using the `cameraPR` function without a fold change threshold. Hallmark gene set collections were obtained from the Molecular Signature Database using the msigdbr package (v7.5.1)

The INTERFEROME database (http://www.interferome.org/interferome/home.jspx) was used to identify and annotate previously defined IRGs among the significantly expressed genes detected, using the following parameters: Species = *Homo sapiens*, Treatment time/concentration = Any; and fold change 2 up/down). The database contains manually curated datasets for type-I, II, and III IRGs with detailed annotation and search capabilities for a comprehensive interpretation of results (36).

The CiiiDER software was used with the human GRCh38.94 genome and the 2020 JASPAR core non-redundant vertebrate matrices to predict transcription factor binding sites (TFBS) and perform TFBS enrichment analysis in the promoter regions of differentially expressed genes (108). Promoter regions for the differentially expressed genes spanning 1,500 bases upstream and 500 bases downstream from the predicted transcriptional start site were used for the analysis. The Match algorithm (109) was used to predict TFBS; the significance of over and underrepresented TFs was calculated using Fisher’s exact test (p < 0.01 and test statistic > 0).

### ChIP-seq analysis

Publicly available, non-overlapping ChIP-seq datasets were extracted from the ENCODE database (https://www.encodeproject.org/) (68). The Bedtools application was used to calculate the overlap between bed files from ENCODE and the outputs from CiiiDER analysis (110).

## Data availability

R1 and R2 FASTQ files were de-multiplexed using cutadapt v3.3 (with error rate 0 and action none) for deposition in ENA with accession PRJEB121415.

## Code availability

Code used during data analysis is available through Monash Bridges (https://doi.org/10.26180/33091607).

## Acknowledgements

This work was supported by the National Health and Medical Research Council (NHMRC) through project grants 1125704 (to G.W.M. and P.R.G.), 1160838, and 1079211 (to G.W.M.); the Australian Research Council through grants DP210100998 (to P.R.G. and G.W.M) and DP150102569 (to G.W.M.); and a Grimwade Fellowship from the Meigunyah Fund awarded to G.W.M.

---

**Supplementary Figure 1.**
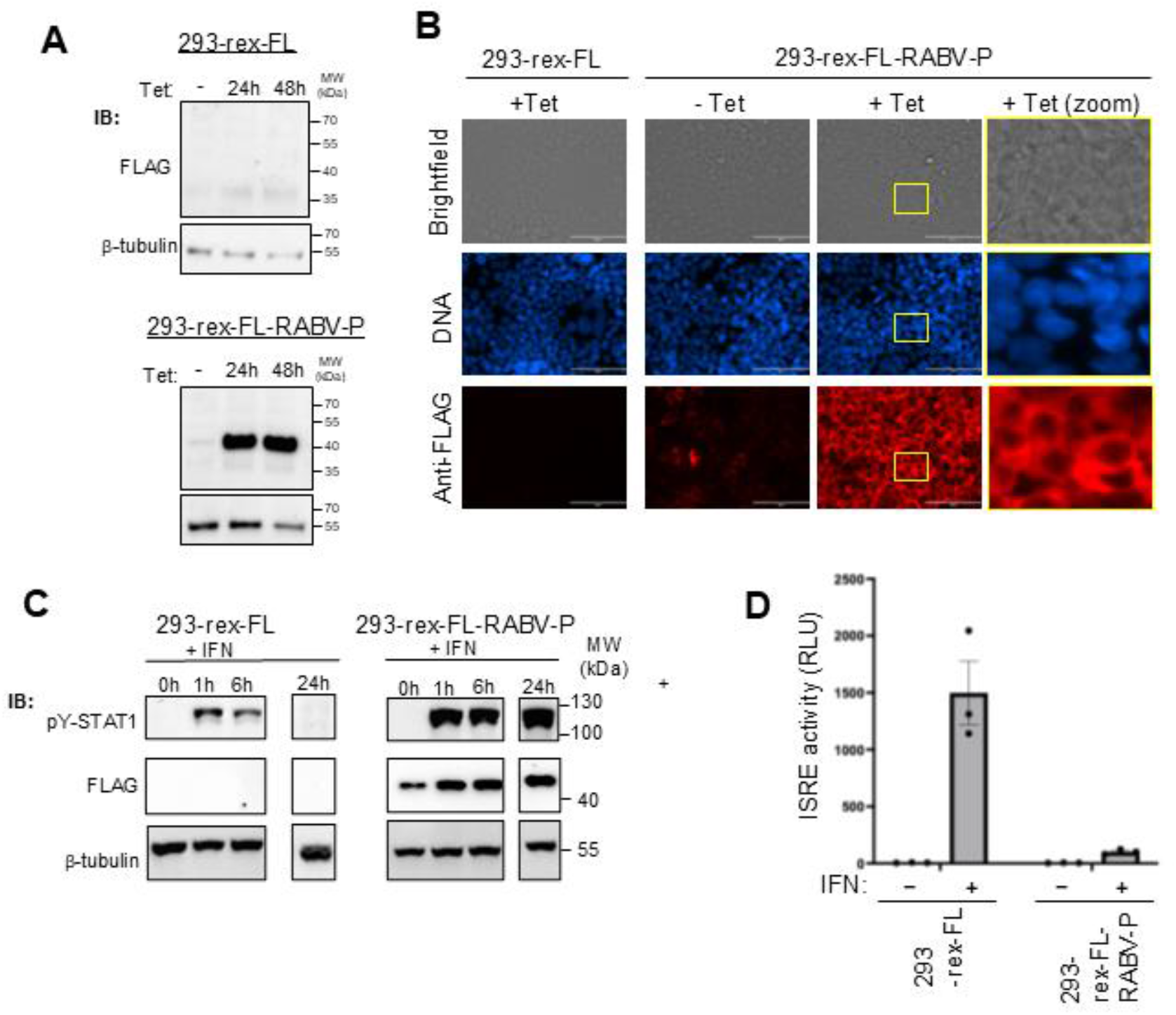
Validation of inducible RABV-P protein expression and function in Flp-In^TM^ T-Rex™-293 cells. (A) IB analysis of RABV-P protein expression in 293-rex-FL and 293-rex-FL-RABV-P cells. Cells were treated with 1 µg/ml tetracycline (Tet) for 24 or 48 h before lysis and IB blot analysis of lysates with the indicated antibodies. (B) Representative images captured using an EVOS fluorescence microscope. 293-rex-FL and 293-rex-FL-RABV-P cells were imaged with or without Tet induction (1 μg/ml, 24 h). Panels show brightfield, nuclear staining with Hoechst 33342 (DNA, blue), and anti-FLAG immunofluorescence (red). Insets show regions enlarged in zoom panels. (C) IB blot analysis of pY-STAT1 in Tet-induced 293-rex-FL and 293-rex-FL-RABV-P cells following IFN-α treatment for the indicated times (irrelevant lanes removed). (D) Tet-induced RABV-P protein inhibits type-I IFN-dependent reporter gene expression. 293-rex-FL and 293-rex-FL-RABV-P cells were transfected with the pISRE-luc plasmid in which firefly luciferase expression is under the control of an IFN-stimulated response element (ISRE)-containing promoter and pRL-TK plasmid (which expresses Renilla luciferase constitutively) for 6 h prior to Tet induction. Cells were then treated with IFN-α (1000 U/ml) or left untreated for 16 h. Relative luminescence units (RLU) for ISRE activity were calculated by normalizing firefly luciferase activity to *Renilla* luciferase activity.

